# Finite-size transitions in complex membranes

**DOI:** 10.1101/2020.11.05.370189

**Authors:** M. Girard, T. Bereau

## Abstract

The lipid raft hypothesis postulates that cell membranes possess some degree of lateral organization. The last decade has seen a large amount of experimental evidence for rafts. Yet, the underlying mechanism remains elusive. One hypothesis that supports rafts relies on the membrane to lie near a critical point. While supported by experimental evidence, the role of regulation is unclear. Using both a lattice model and molecular dynamics simulations, we show that lipid regulation of a many-component membrane can lead to critical behavior over a large temperature range. Across this range, the membrane displays a critical composition due to finite-size effects. This mechanism provides a rationale as to how cells tune their composition without the need for specific sensing mechanisms. It is robust and reproduces important experimentally verified biological trends: membrane-demixing temperature closely follows cell growth temperature, and the composition evolves along a critical manifold. The simplicity of the mechanism provides a strong argument in favor of the critical membrane hypothesis.

**SIGNIFICANCE:** We show that biological regulation of a large amount of phospholipids in membranes naturally leads to a critical composition for finite-size systems. This suggests that regulating a system near a critical point is trivial for cells. These effects vanish logarithmically and therefore can be present in micron-sized systems.

## INTRODUCTION

Lipid rafts—the lateral organization of cell membranes in lipid domains—were postulated decades ago (1, 2). Raft formation would have advantages for biological systems, for instance concentration of reactants and proteins or local regulation of membrane properties, and there is sufficient experimental evidence that some sort of lipid heterogeneity occurs in biological membranes. Yet, many of the controversies remain unsolved: composition of rafts cannot be accurately determined since extraction processes influence raft compositions measured, and while rafts are generally associated with elevated cholesterol, some mass spectroscopy experiments on cells are not consistent with this (3–6); and a number of experiments raise doubts on the existence of rafts (7, 8). The very physical origin of rafts is still up to debate, and many physical mechanisms have been brought forward to explain them (9).

Lipid rafts and associated lipodomes are usually investigated through giant plasma membrane vesicles (GPMV), vesicles extracted from plasma membranes that retain most of their lipid and protein composition (10, 11). When these are cooled by 15 – 30 K below cell temperature, they demix into two distinct phases, which contain slightly different lipid and protein compositions (12, 13). There is experimental evidence showing that the demixing process is critical, and belongs to the Ising universality class (14, 15) (which includes usual two-phase critical points). Proximity to a critical point could yield advantages for cells: high heat capacity and chemical susceptibility — how strongly composition reacts to environmental changes, and could contribute to protein sorting on the membrane through long-range interactions induced by lipids (16).

However, this doesn’t appear compatible with regulation mechanisms. Regulating the composition close to a critical point would require a delicate tuning of all membrane components. This would entail elaborate protein sensing and feedback mechanisms in the membrane. Yet, phospholipid composition is mainly regulated in the endoplasmic reticulum, with no evidence of any mechanism carefully maintaining the composition near a critical point. Furthermore, the lipid composition varies not only with cell type, but also over time for each cell. This indicates that membrane homeostasis does not maintain precisely maintain composition, but rather that composition drifts along a critical manifold. This appears at odd with normal a normal composition-sensing feedback mechanism.

Relations between membrane composition, regulation, and properties are complex and possess many degrees of freedom. For this reason, membrane behavior is often investigated using ternary membranes, a mixture of saturated (usu-ally di-palmitoyl phosphatidylcholine (DPPC)), and unsaturated (usually di-oleyl phosphatidylcholine (DOPC)) lipids along with cholesterol. These membranes can possess a critical point and are used to approximate cellular behavior, as they can demix into two phases, with distinct composition: liquid ordered (DPPC and cholesterol rich) and liquid disordered (DOPC rich). However, the chemical contrast between the two phases, and their nematic order parameter, is much greater than what is found in GPMV. It is unclear to what extent they represent biological membranes, as simulations using biological-like composition tend to show more complex behavior (17–19).

While ternary membranes are well understood, phase behavior of *N*-component mixtures is largely unexplored. Part of the reason is the curse of dimensionality, as every new chemical species adds a dimension to the problem, which rapidly becomes untractable. We have recently established a framework to link lipid regulation to membrane composition, based on chemical potentials between lipids (20). To make computation tractable, we introduced the equal-binding approximation, which postulates that all lipid chemical potentials are equal. This approximation assumes that proteins involved in the Lands’ cycle are non-specific to different phospholipids. Nevertheless, this model successfully reproduced various biological trends, in particular the cholesterol-lipid saturation correlation. In regular solution models, the equal-binding approximation drives the system towards the critical point, if it exists, as equal chemical potentials imply a minimization of the free energy with respect to composition. One would expect this to extend to simple model membranes, so that lipid composition would tend to settle in the vicinity of critical points, provided the temperature is higher than the critical demixing temperature. In model membranes with 2 components and cholesterol, there are clear regions of phase separation, with critical points (19, 21). Naively, for large values of *N*, one would then expect a critical point to exist somewhere in chemical space as cholesterol concentration is changed However, using *N* = 16-component membranes, our model did not show any critical behavior features. This suggests that there are discrepancies in behavior between commonly used simple *N* = 2 membranes and complex *N* = 16 membranes, This naturally poses the follow-up questions: at which value of *N* does a membrane become “complex”, and what are its characteristics?

Here, we propose a coherent framework to study membrane critical behavior together with lipid regulation. Studying critical behavior typically requires sampling large systems over significant timescales, for instance to perform finite-size scaling analysis or overcome critical slowdown; therefore lattice-based models are often preferred due to lower computational requirements. Consequently, we use a lattice model to simulate regulation in membranes and establish a series of trends. To do so, we first extend the Ising-model to *N* states, while conserving the underlying symmetry, and therefore the 2D Ising universality class. We show that, for sufficiently large *N* and under a transition temperature *T*_fs_, finite-size effects in the disordered phase lead to two distinct statistical ensembles. Averaging over the lattice—because of its instantaneous composition—is not equivalent to averaging over the regulated ensemble, due to poor sampling of the lipid distribution.

The implications of the lattice-based models are significant: while living cell membranes are implemented here by the regulated ensemble, the frozen-configuration lattice ensemble emulate GPMV. Our results highlight that the two operate in different thermodynamic ensembles, due to finite-size effects. These finite-size effects are associated with a large value of the correlation length in the regulated ensemble. Critically, the effects persist over a wide range of temperatures. Our results therefore differ from critical behavior, which would force the cell to operate in a narrow range of compositions and temperatures.

GPMV experiments can be emulated by a lattice ensemble with a fixed composition taken from a regulated ensemble at temperature *T*_0_. Biologically, *T*_0_ can be associated with the temperature at which the cells were grown. The observed demixing temperature *T*_m_ in the lattice ensemble is a function of *T*_0_. In particular, for *T*_0_ < *T*_fs_, we observe the relation *T*_0_ ≈ *T*_m_. More importantly, this implies that regulating cell temperature to be close to a critical demixing point is trivial in the finite-size regime, and that chemical composition can fluctuate while staying close to the critical demixing.

We validate our extended Ising model with coarse-grained molecular dynamics simulations, using 122 lipid species. The picture prevails overall, confirming the validity of our lattice-based model. The increased level of resolution naturally brings about more features. In particular, the distribution of lipids—how many lipids of each type is present in the regulated ensemble—is strongly affected by molecular interactions not well captured in the lattice model. This in turn affects the effective number of chemical species present. This distribution can be altered by changing chemical potentials or varying cholesterol. Small amounts of the latter (≈ 5%) rapidly increases *T*_fs_.

## METHODS

### Lattice-based model

We model membranes by an *N* state Ising-like model on a *L* × *L* lattice. The Hamiltonian 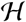 preserves the *Z*_2_ symmetry of the Ising model, such that 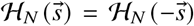. There is no Potts rotational symmetry, and unlike in the Blume-Capel model, all ground states are equivalent. Each site in the lattice takes a chemical state *s_i_* = {−1, −1 + 2/(*N* – 1),…, 1}. This represents a chemical affinity ladder, where lipids interacts with energy only dependent on their distance on this ladder. The energy of the system is given by

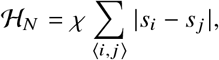

where 〈*i, j*〉 indicates all nearest-neighbor pairs *i, j*, and *χ* is an energy scale representing chemical mismatch between least and most saturated lipids. For *N* = 2, this is exactly the usual 2D Ising model. The chemical state *s* is taken to represent both headgroup affinity and acyl-tail unstaturation.

The limit *N* → ∞ can be taken by letting *s_i_* ∈ [−1, 1] vary continuously. Biologically, this represents a system with an infinite amount of lipid species with continuously increasing chemical mismatch.

We consider both non-conserved and conserved order parameter simulations, corresponding to single-site exchange and two-site swaps, respectively. In order to reproduce GPMV experiments, where chemical compositions are fixed, we impose an additional constraint on conserved order parameter simulations: the overall distribution of spins (e.g., number of spins with *s* = 1) is conserved. As shown later, this crucial aspect leads to two non-equivalent ensembles. Systems are considered equilibrated after 2^20^ Monte Carlo (MC) steps. The number of exchanges per MC step is 4*L*^2^ for non-conserved parameters, and *L*^2^log_2_(*L*)/4 for conserved order parameters, with non-local swaps (see SI for details). We make use of replica exchange using 2048 replicas, and attempt a swap on all replicas every 4 MC step. After equilibration, statistics are gathered every 4 MC steps over 2^26^ steps for non-conserved parameters and 2^19^ steps for conserved order parameters.

### Molecular dynamics simulation

Molecular dynamics simulations are run using the HOOMD-Blue molecular dynamics engine (22, 23) using the MARTINI coarse-grained model (24). Lipids are run in the semigrand canonical ensemble, where their chemical state can freely change. Chemical regulation includes (*i*) headgroup, either phosphatidylcholine (PC) or phosphatidylethanolamine (PE), (*ii*) acyl-tail length, varies via a ghost bead (see SI for details), and (*iii*) acyl-tail unsaturation.

We consider a restricted set of acyl tails. The sn-1 acyl tail has 0 to 2 unsaturations and has 4 beads (16-18 carbons). The sn-2 acyl tail has 0 to 5 unsaturations and 4 - 5 beads (16 – 22 carbons). We restrict the sn-2 acyl tail to have at least as many unsaturations as sn-1, long tails (≥ 20 carbons) to be highly unsaturated (≥ 3), and unsaturations placed consecutively, Considering headgroups, unsaturation and acyl tail length, this yields 122 lipid species in the bilayer. Our bilayers have 800 lipids per leaflet. The full list of acyl tails is available in the SI.

## RESULTS

### Lattice-based model

We will describe membrane simulations that are both regulated and those that have a fixed composition, involving non-conserved and conserved order parameters, respectively In the lattice, any lipid can be exchanged for another lipid type. This is meant to proxy the regulation mechanism. Fixed-composition membranes aim at emulating extracts from GPMV experiments, which probe the phase behavior of a fixed composition taken from a regulated membrane.

### Regulated composition

In this section, we first derive critical properties of the finite *N* model and show appearance of strong finite-size effects at large values of *N*. We show that critical temperature vanishes with *N* as 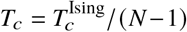, while leaving finite-size effects untouched, and derive conditions for the onset temperature *T*_fs_.

Since the system conserves the *Z*_2_ symmetry, all critical behavior is within the Ising universality class. The first major difference with Ising arises in the difference between the symmetries of odd and even *N* groups. For odd *N*, the element *s_i_* = 0 is allowed and it always maps to itself under the *Z*_2_ symmetry. This causes criticality to vanish for all odd *N* values (see SI for details and further discussion). As *N* increases, the system becomes dominated by finite-size effects, characterized by system-spanning correlation lengths, and the distinction between odd and even *N* vanishes.

Another significant departure from Ising behavior lies in how the correlation length peaks. Fixing *L* but increasing *N*, the peak becomes wider (Fig. 1A). This presents a significant departure from a 2D Ising model, because the system here shows critical behavior over a *range* of temperatures. This behavior manifests itself in the disordered region (*T* > *T*_c_) and up to a temperature *T*_fs_, the latter being independent of *N*. This behavior persists as *N* → ∞, as shown in Fig. 1B. Introducing increasingly more states in the model opens up the temperature span between *T*_c_ and *T*_fs_. Importantly, there is no phase separation at *T*_c_ < *T* < *T*_fs_, the system exhibits instead critical composition. We will show below that this stems from finite-size effects. However, the observations here are not the usual finite-size effects that appear in the vicinity of critical points, because criticality disappears (*T*_c_ → 0) in the limit *N* → ∞.

**Figure 1:**
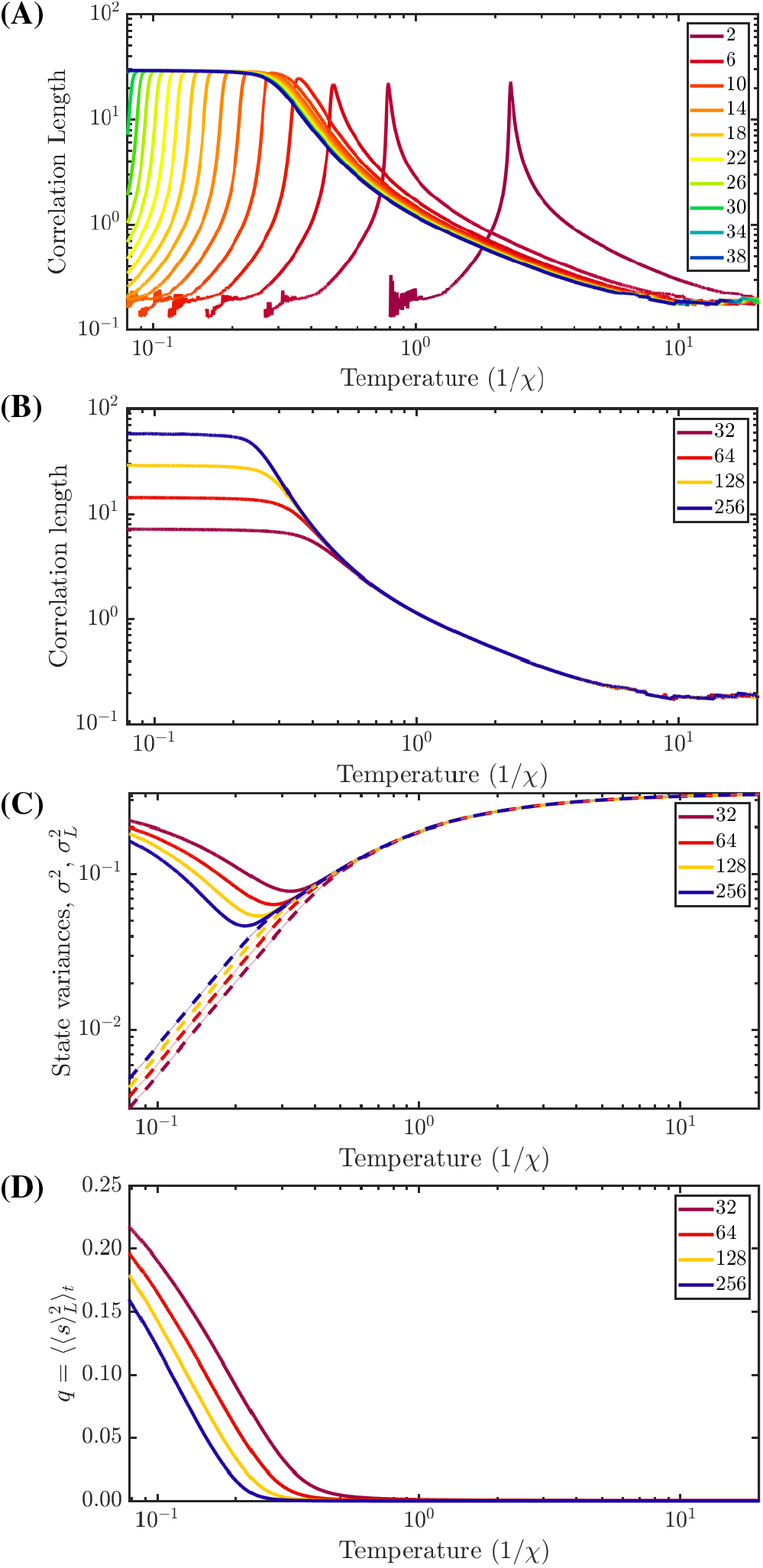
(A) Correlation length of systems of even *N* for *N* ∈ [2,40] for *L* = 128. (B) Correlation length of *N* = ∞ systems for various *L*. (C) Ensemble variance (*σ*^2^, full lines) and instantaneous lattice variance (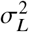, dashed lines) across temperatures for the continuous (*N* = ∞) case for various system sizes. In the low temperature regime, *σ*^2^ = 1/3 and 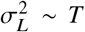, while at high temperatures 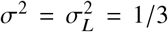. (D) Edwards-Anderson order parameter for *N* = ∞.

To garner insight into this behavior, we characterize the lipid states in the lattice model by their distributions, and label the time-averaged distribution 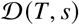 and the instantaneous lattice-average 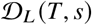. We denote time average as 〈·〉_*t*_ and lattice (spatial) average as 〈·〉_*L*_. The two distributions are related by 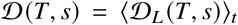. In disordered phases, the first moment 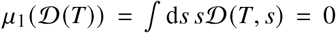, while 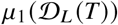 can take non-zero values. We characterize these two distributions by their second moment (variance): 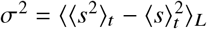 and 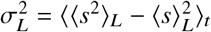.

As shown in Fig. 1C, for *T* < *T*_fs_, we find 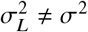. Under *T*_fs_, a single realisation of the lattice average cannot correctly sample the ensemble average. The amount of states sampled is dependent on the size of the lattice, and this onset temperature is dependent on *L*. This is reminiscent of quenched disorder in glasses, where averages need to be taken over many realizations of disorder (see (25) and references within). In our regulated picture, the many realizations of disorder are provided by the non-conserved parameter ensemble, and accordingly, the Edwards-Anderson order parameter 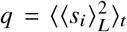 also takes non-zero values for *T* < *T*_fs_. This translates to a large constant value of the correlation length over a wide temperature range. For infinite systems, the correlation length likely corresponds to the wavelength of an oscillation of the local realizations, similar to a Goldstone mode. We note that this regime is destroyed by any ordering of the system, and therefore is only found in the temperature range *T_c_* < *T* < *T*_fs_.

Equivalently, This finite-size transition can be seen as arising from equivalent distributions in the system. Effectively, if the tails of 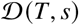 are poorly sampled, then 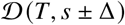 has the same energy and the two distributions are equivalent. The tails are poorly sampled when

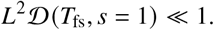

Based on our simulations (see Fig 1), the instantaneous variance follows *σ_L_* ~ *T* at low temperatures. We postulate that at low temperature, the tails are exponentially suppressed, which is justified by measuring the instantaneous distribution (see SI). In such cases, the condition can be expressed as

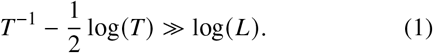

Using Eq. 1 as a basis and computing the flatness onset for *N* = ∞, We find the finite-size onset can be fitted to (see SI):

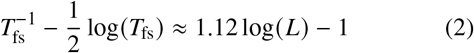

### Fixed composition

We emulate GPMV-experiment extracts by first equilibrating membranes with *L* = 128 using regulation (i.e., a nonconserved order parameter). We then take a composition simulated at temperature *T*_0_ and freeze the composition (i.e., conserved order parameter). In the regime *σ* ≠ *σ_L_*, the two ensembles are not equivalent, as the constraint on the conserved order parameter enforces 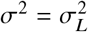. This can be likened to a simulation of a random distribution of chemical states, with a variance constrained to 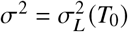. For *T* < *T*_0_, there exists a temperature *T*_2_ = *T_m_* at which the lipids critically phase separate into two distinct phases, which we associate with the experimentally observed GPMV demixing tempeature. Phase behavior is unfortunately complicated by a cascade of phase transitions into 2,3,4,…, phases with distinct demixing temperatures *T*_2_ > *T*_3_ > *T*_4_ > …, which limits our ability to precisely compute *T_m_* (see SI for details). Nevertheless, we characterize it by making use of Gibbs ensemble and a modified Edwards-Anderson parameter, which yields a sufficiently good approximation of the transition temperature. Under *T_m_*, the resulting two phases have similar compositions, characterized by the ratio of variances 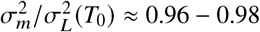.

As shown in Fig. 2, the demixing transition is a function of *T*_0_. For initial temperatures *T*_0_ < *T*_fs_, we find the ratio *ν*(*L*) = *T*_0_/*T_m_* takes a constant value. This implies that the reduced temperature *t* = (*T*_0_ – *T_m_*)/*T_m_* = *υ* – 1 is kept constant by cells. This ratio *υ*(*L*) exhibits a power-law dependence on system size, such that 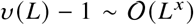, which is expected: near critical points, the correlation length, *ξ*, varies as *ξ* ~ |*t*|^−*ν*^, and under the assumption that *L* ~ *ξ*, we find (*υ* – 1) ~ *L*^−*ν*^. However, fitting the data of Fig. 2 yields (*υ* – 1) ~ *L*^−0.63^. This discrepancy may stem from additional apparent power-laws: 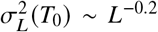 and *T_m_*|*σ*^2^ ~ *L*^−0.13^, whose origin remains elusive.

**Figure 2:**
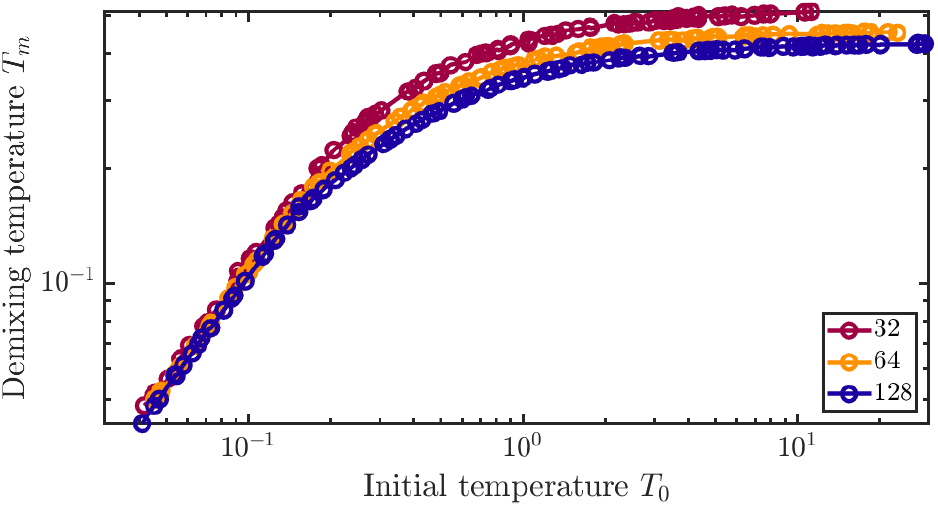
Observed demixing temperature *T_m_* versus initial lattice temperature *T*_0_ for different values of *L*. For temperatures 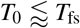, the relation *T*_0_ ≈ *υ*(*L*)*T_m_* is verified.

### Molecular dynamics simulations

Characterization of lipid composition in the membrane is more challenging from coarse-grained molecular dynamics than it is from lattice models. The finite-size transition still manifests itself through a nearly constant correlation length with respect to temperature for *T* < *T*_fs_ (Fig. 3). Correlation lengths measured are on the sub-nanometer lengthscale, similar to our previous simulations (20), and fall below measurable levels at *T* > *T*_fs_, which is in agreement with the lattice-based model.

**Figure 3:**
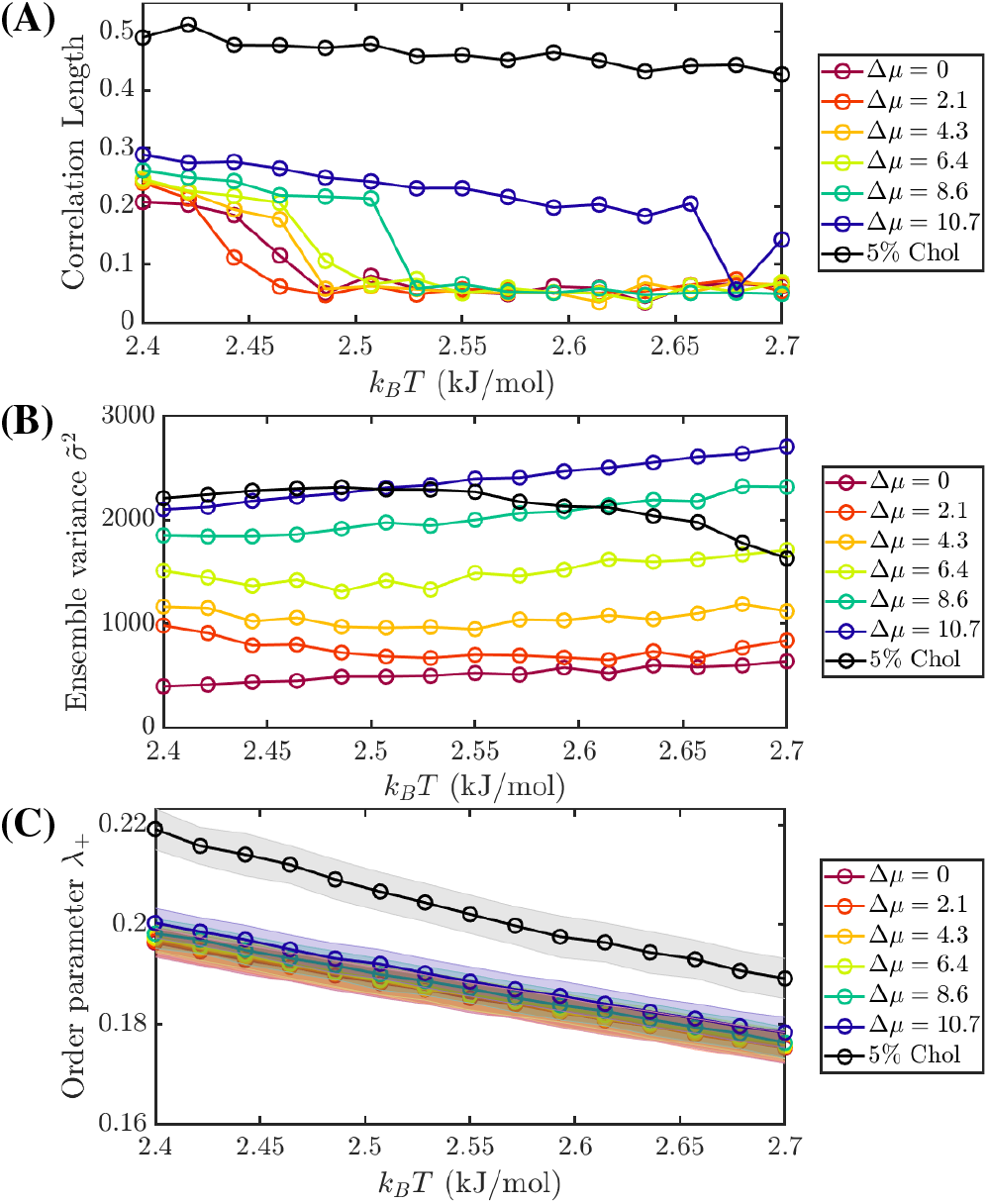
Properties of membranes in molecular dynamics for various values of chemical potential differences between short and long lipids, or when cholesterol is added. (A) Correlation length of membranes, (B) Ensemble variance obtained through histogram approximations, and (C) Nematic order parameter. Shaded bar indicate standard deviation. Chemical potentials are given in kJ/mol

Interactions in chemically-realistic membranes are more complex than simple lattice models. Interactions can take place over longer range than what the lattice model assumes, e.g., by coupling with nematic order. Additionally, the well-defined chemical coordinate *s* is lost and instead replaced by the identity and interactions of each lipid in the system. To characterize the transition, ensemble and instantaneous volume variances, 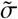 and 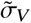, can be approximated by histogram-based methods (see SI). However, due to the lack of a suitable projection onto *s*, these values cannot be used for quantitative comparison.

In our simulations, lipid length is varied by means of a “ghost” bead. Long chains gain more free energy than short chains upon insertion in membrane, which favor presence of long chains in the membrane. Due to the restrictions we placed on lipid species, this leads to a lipodome dominated by C20:3 fatty acids (see SI). In turn, this translate into other lipid species having very low populations or being completely absent. The dominance of C20:3 fatty acids can be reduced by tuning the chemical potential between short and long lipids (Δ*μ*_*C*16–*C*20_). Additionally, cholesterol can be added in the simulation. Due to its preference for saturated lipids, this strongly favours short lipids. Both of these changes translate into increases of *T*_fs_ observed in simulations, interestingly without alterating the finite-size behavior observed (Fig. 3).

We propose here two potential explanations for this behavior. First, there is a change of effective range of the chemical coordinate *s*, which we can visualize by changes in 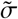 (Fig 3). In the lattice model, the range of *s* is implicitly included in the definition of *χ*, therefore increasing the range of *s* is equivalent to decreasing temperature. Second, the nematic order increases with Δ*μ*_C16–*C*20_ and particularly with cholesterol (Fig 3C). Increases in nematic order can influence lipid-lipid interactions and lead to an effective increase of *χ*.

## DISCUSSION

### Regulation near criticality

The simple lattice-based model demonstrates the possibility to regulate a complex *N*-component system near criticality, provided *N* is sufficiently large and molecules suitably distributed in chemical space. It does not require any particular feature of the system, except that there exists a cycling mechanism that can transform the chemical state of lipids. This process has two hallmarks associated with it. First, freezing the composition at cell-growth temperature *T*_0_ < *T*_fs_ and changing temperature yields an observed demixing temperature that depends on *T*_0_. Second, instantaneous composition drifts in chemical space along a critical manifold. Both of these are known biological features of cell membranes (26), and have been part of the lipid-raft debates. In regulated ensembles, they arise naturally.

However, this model does not replicate the constant *T*_0_ – *T_m_* ≈ 15 – 30 K observed in GPMV experiments. Rather, we predict *T*_0_ ≈ *T_m_* for micron-sized systems. A potential explanation for this discrepancy stems from the membrane cytoskeleton, a network of actin fibers. Transmembrane proteins are located along this network and hinder diffusion as well as phase separation and would prevent the large clusters predicted in Fig. 1B (27). The typical length scale of this network is in the range of 40 – 230 nm (28). A system representative of this size (*L* ≈ 64 – 256) would exhibit *ν* ≈ 1.03-1.1, which translates to *T*_0_ – *T_m_* ≈ 7 – 30 K, in-line with experimental observations.

### Density of chemical states

The lattice model here uses an equal-binding approximation (Δ*μ* = 0) in conjunction with linearly spaced lipids on the chemical *s* space. The latter implies that the density of chemical states, Ω(*s*) — how many chemical species exist within the interval (*s* – *ε, s*+*ε*) — is constant. We note that the chemical potential of specie *s* can be related to its local density of states by *μ*(*s*) = *k_B_T* log(Ω(*s*)). How the system behavior depends on these approximations remains unknown. Simulations with a normal distribution 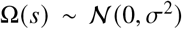 yield similar results. Similarly, molecular dynamics simulations suggest that the mechanism is unaffected by changes in chemical potential values, up to a change in *T*_fs_. However, whether *υ*(*L*) changes with the density of states is yet unexplored.

### Cholesterol, proteins, and rafts

Our model does not predict raft formation. A potential gateway for raft formation is through unregulated components — components that are not directly regulated by chemical potential differences but rather have prescribed fractions. Every protein possesses a unique spatial distribution of lipid binding domains, the so-called protein fingerprints (29), which influences local lipid composition. In the vicinity of critical points, components immersed in the critical fluid experience long-range interactions (30). This force, called critical Casimir interactions (31) has been proposed as a membrane-protein organization pathway (16). However, in the context of lipid rafts, for low temperatures *T* ≪ *T*_fs_, *σ_L_* ≪ *σ* and the mean state can dynamically change, and in turn, affect the lipid-protein affinities.

Membrane behavior with 5% cholesterol appears at first glance to have transition temperatures higher than *k*_B_*T* = 2.7 kJ/mol. However, the significantly higher value of correlation length (≈ 0.5 nm) than for membranes without cholesterol (≈ 0.20 – 0.3 nm) suggests that other processes are at play. This behavior is reminiscent of colloids under critical Casimir forces, where the structure factor at *k* = 0 shows colloid fraction dependence (32) and will need further investigation. Whether similar behavior is observed for other unregulated components, e.g., sphingolipids and membrane proteins, is also unclear.

For membranes, the cycling process is done by the Lands’ cycle. In principle, this regulation mechanism is not limited to two-dimensional systems (membranes), and should be applicable to other regulated systems, such as the cytoplasm. In this context, lipids would be analogous to the sea of disordered proteins, the Lands’ cycle with poly-ubiquitination, and lipid rafts with membraneless organelles. However, it is unclear in this picture how the membraneless organelles survive since they would be dissolved by poly-ubiquitination.

### Regulated ensembles and GPMV experiments

A particular feature of chemical composition below *T*_fs_ is the distinction between volume (or lattice) and ensemble averages. Experiments that measure lipidome of GPMV typically perform measurements over multiple different GPMV, which is essentially an ensemble average. While this can provide insights into cell behavior, the ensemble composition can be quite different (especially at low temperatures) than the membrane composition of a single cell. In principle, the mechanism proposed here can be verified by experimental sampling of 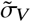 and 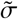, and a suitable projection to a chemical coordinate. However, this also requires extensive use of single-cell lipidomics (33, 34), an emerging technique. Furthermore, experiments can be further complicated by cell-to-cell variability, in particular of non-regulated components, e.g., membrane proteins.

## CONCLUSIONS

We presented here a study of *N*-component membranes allowing for lipid regulation, using both a lattice-based approach and molecular dynamics. For properly distributed chemical species and sufficiently large *N*, the system exhibits a finite-size temperature, which stems from differences between lattice and time averaging. Under this temperature, composition drifts near a critical manifold. Unlike usual critical behavior, system-spanning correlation lengths are observed in a wide temperature range. Furthermore, molecular dynamics simulation suggests that this mechanism is robust with perturbations of chemical potentials in the system. Taken together, these results show that composition regulation near criticality is trivial for cells and that it needs no sensing mechanism.

These results also open an important new question: how do proteins behave on this membrane ? Studies on critical Casimir interactions tend to model colloids with uniform surfaces in a critical solvent and show rich behavior. However, biological membranes contain many different proteins, each with a unique fingerprint, and the solvent composition changes with protein composition. How this affects aggregation and their ability to form raft-like domains is an exciting question for biological organization. We believe that the regulated ensemble picture can help elucidate some of these questions.

## AUTHOR CONTRIBUTIONS

M.G. and T.B. designed the research. M.G. carried out all simulations, analyzed the data. M.G. and T.B. wrote the article.

## ACKNOWLEDGMENTS

We acknowledge Burkhard Dünweg for many insightful discussions and Kurt Kremer, Thomas Shaw, Sarah Veatch, Benjamin Machta and Isabella Graf for a critical reading of this manuscript.

We acknowledge financial support from the Alexander von Humboldt Sitftung (AvH) and the Deutsche Forschungsgemeinschaft (DFG) and computational ressources from the Max-Planck computing and data facilities (MPCDF).

## SOFTWARE

Lattice based models were run using custom in-house software. Molecular dynamics simulations were run in the HOOMD-Blue molecular dynamics engine (22, 23) with a custom in-house semi-grand canonical plugin. Initial configurations were assembled using the hoobas molecular builder (35). Molecular dynamics simulations use a slightly modified MARTINI force-field (24) (see SI). Visualization was done using the Ovito package (36). Analysis is done using in-house code.

## SUPPLEMENTARY MATERIAL

An online supplement to this article can be found by visiting BJ Online at http://www.biophysj.org.

